# Vps34 PI 3-kinase controls thyroid hormone production by regulating thyroglobulin iodination, lysosomal proteolysis and tissue homeostasis

**DOI:** 10.1101/580142

**Authors:** Giuseppina Grieco, Tongsong Wang, Ophélie Delcorte, Catherine Spourquet, Virginie Janssens, Aurélie Strickaert, Héloïse P. Gaide Chevronnay, Xiao-Hui Liao, Benoît Bilanges, Samuel Refetoff, Bart Vanhaesebroeck, Carine Maenhaut, Pierre J. Courtoy, Christophe E. Pierreux

## Abstract

**BACKGROUND:** The production of thyroid hormones (T_3_, T_4_) depends on the organization of the thyroid in follicles, which are lined by a monolayer of thyrocytes with strict apico-basal polarity. This polarization supports vectorial transport of thyroglobulin for storage into, and recapture from, the colloid. It also allows selective addressing of channels, transporters, ion pumps and enzymes to their appropriate basolateral (NIS, SLC26A7 and Na^+^/K^+^-ATPase) or apical membrane domain (Anoctamin, SLC26A4, DUOX2, DUOXA2 and TPO). How these actors of T_3_/T_4_ synthesis reach their final destination remains poorly understood. The PI 3-kinase (PI3K) isoform Vps34/PIK3C3 is now recognized as a main component in the general control of vesicular trafficking and of cell homeostasis via the regulation of endosomal trafficking and autophagy. We recently reported that conditional Vps34 inactivation in proximal tubular cells in the kidney prevents normal addressing of apical membrane proteins and causes abortive macroautophagy.

**METHODS:** Vps34 was inactivated using a Pax8-driven Cre recombinase system. The impact of Vps34 inactivation in thyrocytes was analyzed by histological, immunolocalization and mRNA expression profiling. Thyroid hormone synthesis was assayed by ^125^I injection and serum plasma analysis.

**RESULTS:** Vps34^cKO^ mice were born at the expected Mendelian ratio and showed normal growth until postnatal day 14, then stopped growing and died at around 1 month of age. We therefore analyzed thyroid Vps34^cKO^ at postnatal day 14. We found that loss of Vps34 in thyrocytes causes: (i) disorganization of thyroid parenchyma, with abnormal thyrocyte and follicular shape and reduced PAS^+^ colloidal spaces; (ii) severe non-compensated hypothyroidism with extremely low T_4_ levels (0.75 ± 0.62 μg/dL) and huge TSH plasma levels (19,300 ± 10,500 mU/L); (iii) impaired ^125^I organification at comparable uptake and frequent occurrence of follicles with luminal thyroglobulin but non-detectable T4-bearing thyroglobulin; (iv) intense signal in thyrocytes for the lysosomal membrane marker, LAMP-1, as well as thyroglobulin and the autophagy marker, p62, indicating defective lysosomal proteolysis, and (v) presence of macrophages in the colloidal space.

**CONCLUSIONS:** We conclude that Vps34 is crucial for thyroid hormonogenesis, at least by controlling epithelial organization, Tg iodination as well as proteolytic T_3_/T_4_ excision in lysosomes.

## Introduction

The main function of the thyroid gland is to produce the hormones, T_3_/T_4_ and calcitonin, which are essential for the regulation of metabolic processes (1, 2). The production of T_3_ and T_4_ depends on the correct tissue organization of thyroid epithelial cells, the thyrocytes, into functional and independent units, the follicles. These are composed of a single layer of polarized thyrocytes that form a spherical structure delineating an internal space or lumen where the thyrocyte secretory product, thyroglobulin, is stored in a colloidal form, thus called the colloid lumen. As thyrocytes communicate via gap junctions, each follicle functions as an integrated unit. Thyroid hormone synthesis depends on apico-basal cell polarity that allows the specific localization of channels, transporters, pumps and enzymes at the appropriate membrane domains. Iodide from the bloodstream freely traverses the fenestrated endothelium of the thyroid capillaries and is taken up into thyrocytes via the basolaterally-localized Na^+^/I^−^ symporter (NIS), thanks to a Na^+^ gradient generated by the Na^+^/K^+^-ATPase. An alternative basolateral transporter, SLC26A7, has been recently reported to also control iodide uptake, although its role might be indirect (3). Iodide diffuses freely within thyrocytes and is next transported across the apical membrane into the colloid space via apically-localized transporters such as anoctamin or pendrin (SLC26A4). Iodide is then rapidly oxidized into iodine by thyroperoxidase (TPO) located at the apical membrane, in the presence of H_2_O_2_, generated by the DUOX2/DUOXA2 apical membrane complex. Iodine is incorporated into accessible tyrosine residues close to the N- and C-termini of thyroglobulin (Tg), a large protein secreted by thyrocytes into the colloid space. Iodotyrosine rearranges into hormonogenic peptides, which are the direct T_3_ and T_4_ precursors. Thyroid hormone synthesis thus requires basal localization of NIS, SLC26A7 and Na^+^/K^+^-ATPase, apical localization of anoctamin, pendrin, TPO, DUOX2 and DUOXA2, as well as apical delivery of Tg into the colloid lumen and endocytic uptake of iodoTg into thyrocyte lysosomes (1, 2). Specific regulators of thyroid follicular organization have recently been identified (4). Transcriptomic comparison of thyroid FRT cells cultured in 2D monolayers and in 3D spherical follicles indeed revealed involvement of structural and functional cell elements such as adherens and tight junctions (cadherin-16), cytoskeleton proteins, ions channels, proteins involved in differentiation, and components of the trafficking machinery (e.g. myosin-Vb, Rab17) (4).

There is strong evidence that vesicular trafficking is critical for apico-basal polarization and epithelial function (5–7), but the role of vesicular trafficking in thyroid function is incompletely understood. Vps34/*PIK3C3* (also referred to as type III PI3K) has long been recognized as a main actor involved in the general control of endocytic vesicular trafficking (8–13). Vps34/PIK3C3 also plays an important role in epithelial organization in *Drosophila* (14) and in autophagy (12, 15), both in the initiation of autophagosome formation and the progression towards autophagosome-lysosome fusion (16, 17).

We recently inactivated Vps34 in kidney proximal tubular cells (PTC) using Wnt4-Cre and Pax8-Cre (18, 19). Wnt4-Cre driven Vps34 excision caused major perinatal lethality, with abnormalities in kidney cortex development and PTC apical differentiation in the surviving mice. With the Pax8-Cre, which triggers recombination in nephrogenic tubules and in nonvascular components of glomeruli two days later than Wnt4-Cre (20, 21), we observed normal mouse growth until postnatal (P) day 14 (19). Then, body weight levelled off and all pups died at ∼3-5 weeks of age, probably due to kidney failure. Structural and functional studies of Vps34^cKO^ kidneys revealed preserved PTC membrane polarity, but impaired apical membrane protein trafficking, thus causing a general PTC dysfunction known as renal Fanconi-like syndrome, manifested by polyuria and low-molecular weight proteinuria. Vps34^cKO^ also displayed impaired lysosome size/positioning and blocked autophagy, thereby causing cell vacuolization. We concluded that Vps34 is a crucial component of the trafficking machinery necessary for differentiated PTC function and is essential for overall PTC homeostasis (19). Given that Pax8-Cre also triggers recombination in thyrocytes (21), we here investigated the role of Vps34/PIK3C3 in thyroid function and homeostasis.

## Materials and methods

### Mice

Vps34^fl/fl^ mice have been described (22). Pax8-Cre mice were obtained from Dr. M. Busslinger (21). Vps34^fl/fl^ mice were crossed with Pax8-Cre;Vps34^fl/+^ mice to generate conditional targeted excision of Vps34 exon 21 in the thyroid of 25% of offspring (Vps34^cKO^). Non-recombined Vps34^fl/fl^ mice were used as control. Mice were treated according to the NIH Guide for Care and Use of Laboratory Animals, and experiments were approved by the University Animal Welfare Committee, Université Catholique de Louvain (2016/UCL/MD/006 and 2018/UCL/MD/026).

### Plasma, tissue collection and histology

Blood was collected by eye sinus puncture at sacrifice (P14) under irreversible anesthesia by xylazine 2% and ketamine 50 mg/ml (200 lll l/mice i.p.). Thyroid lobes were excised, fixed by immersion in neutral-buffered formaldehyde (4% F) at 4°C under stirring overnight. Samples were paraffin-embedded or equilibrated overnight in 20% sucrose and embedded in Tissue-Tek Optimal Cutting Medium (Sakura Finetek) for cryostat sections.

### TSH and T4 plasma concentrations

Plasma TSH concentrations were measured by a sensitive, heterologous, disequilibrium double-antibody precipitation RIA as described (23). T4 concentration was measured by coated-tube RIA (Siemens Medical Solution Diagnostics, Los Angeles, CA).

### Immunofluorescence

Immunofluorescence was performed on 5 µm-thick frozen sections or on 6 µm-thick paraffin sections (19). Antigen retrieval was promoted in citrate buffer, pH 6.0, at 98°C for 20 min using a Lab Vision Pretreatment Module™ (Thermo Scientific). After permeabilization with PBS/0.3% Triton-X100 for 5 min, non-specific sites were blocked by 1-h incubation in PBS/0.3% Triton-X100 with 10% bovine serum albumin (BSA) and 3% milk, followed by primary antibodies (described in Supplementary Table I) in blocking buffer at 4°C overnight. After extensive washing, sections were incubated with the appropriate AlexaFluor-secondary antibodies in 10% BSA/0.3% Triton-X100 at room temperature for 1 h, extensively washed, mounted with Faramount Aqueous Mounting Medium (Dako) and imaged on a spinning disk confocal microscope using a Plan Apochromat 100x/1.4 Oil DIC objective (Cell Observer Spinning Disk; Zeiss). For whole thyroid section recording, images were acquired using Zeiss Pannoramic P250 slide scanner, stitched and analyzed using Case Viewer software.

### RT-qPCR

Total RNA was extracted from thyroid lobes using TRIzol Reagent (Thermo Scientific), as described (24). Aliquots of 500 ng RNA were reverse-transcribed by M-MLV reverse transcriptase (Invitrogen) with random hexamers, as described (25). Primer sequences used are described in Supplementary Table II. Real-time qPCR was performed as described (25), in presence of 250 nM of specific primers with Kappa SYBR Fast qPCR Master Mix (Kapa Biosystems) on a CFX96 touch real-time PCR Detection System (Bio-Rad). Data were analyzed using the ΔΔCT method, using the geometric mean of □-*Actin* and *Rpl27* as reference genes (26).

### ^125^I uptake and organification

At postnatal day 8 (P8), mothers and litters were fed with an iodine-free diet for 24 h. At P9, pups were injected intraperitoneally with 1 µCi ^125^I (Perkin Elmer) and left for another 24 h before sacrifice and thyroid dissection. Both lobes were collected in 2 mM methimazole (Sigma) and homogenized with a glass Potter. Total thyroid ^125^I was measured with an automatic gamma counter “Wizard^2^” (Perkin Elmer) before protein precipitation using 10% trichloroacetic acid (TCA; Merck). After a single wash in TCA, radioactivity was measured in the protein pellet. Percentage of protein-bound iodide was calculated using the ratio of precipitated cpm/total thyroid cpm (27).

### H_2_O_2_ level measurements

The hydrogen peroxide Assay kit from Abcam (ab102500) was used. At P14, thyroid lobes were collected in ice-cold PBS and homogenized with a glass Potter. Proteins were precipitated with perchloric acid 4 M, and supernatant was neutralized with KCl 3 M and KCl 1 M until pH was comprised between 6.5 and 8. Fifty □l, corresponding to a ninth of a thyroid, was incubated with OxiRed probe and horseradish peroxidase (HRP), according to the instructions form the manufacturer. Fluorescence was measured with a GloMax fluorimeter (Promega, Ex/Em = 535/587 nm).

### Statistical analysis

All statistical analyses were determined by Prism software (GraphPad Software, La Jolla, California, USA). RT-qPCR values were obtained by the ΔΔCT method and are expressed as boxplots with median, 25^th^ and 75^th^ percentiles, and min-max whiskers. Each graph represents the results form a minimum of 8 independent thyroid lobes from at least 3 different litters. Nonparametric statistical tests were used: Mann-Whitney for single comparisons. Differences were considered statistically significant when p<0.05 (*); ** stands for p<0.01; *** for p<0.001.

## Results

### Genetic construction and assessment of Vps34 inactivation

Ubiquitous Vps34 inactivation is embryonically lethal, with embryos dying around embryonic day E8.5 (28). To study tissue-specific roles of Vps34 *in vivo,* we used a mouse line which carries a floxed *Vps34* allele (Vps34^fl^) that allows to conditionally delete the loxP-flanked exon 21 of the *Vps34* gene, which encodes a critical sequence of the lipid kinase domain of Vps34 (22). This approach was designed to allow for the expression of a minimally-truncated, catalytically-inactive Vps34 protein. However, upon conditional expression in megakaryocytes, the level of this truncated Vps34 and its obligatory partner, Vps15, were found to be decreased by 80-90% in the megakaryocyte lineage (thrombocytes), indicating instability of truncated Vps34 protein (22). We will thus hereafter refer to the result of this Vps34 truncation as Vps34 conditional knock-out (Vps34^cKO^).

We crossed homozygous Vps34^fl/fl^ mice with Pax8-Cre;Vps34^fl/+^ mice, which is expected to lead to a tissue-specific (thyroid and kidney) conditional excision of exon 21 from both Vps34 alleles in 25% of the pups, which are further referred to as Vps34^cKO^ mice. Vps34^cKO^ mice were born at the expected Mendelian ratio and showed normal development until postnatal day 14 (P14), then stopped growing and died at around 1 month of age (19). We therefore analyzed the thyroid of Vps34^cKO^ mice at P14. Of note, eye opening, a marker of cerebral maturation was usually delayed till P10.

We first assessed the extent of Vps34 genetic excision in the thyroid by quantification of *Vps34* mRNA using primer that are specific for the total *Vps34* or are exon 21-specific. Compared to control littermate pups, we found a ∼70 % reduction of exon 21-containing *Vps34* mRNA in cKO thyroid extracts. In comparison, we found no significant difference in the abundance of *Vps34* mRNA containing exon 22-24 (Fig. 1A). Assuming that *Vps34* is equally expressed in all thyroid cell types (expressing and non-expressing the Cre recombinase), this ∼70 % decrease suggests that most thyrocytes in Vps34^cKO^ mice had undergone Cre-mediated recombination.

**Figure 1.**
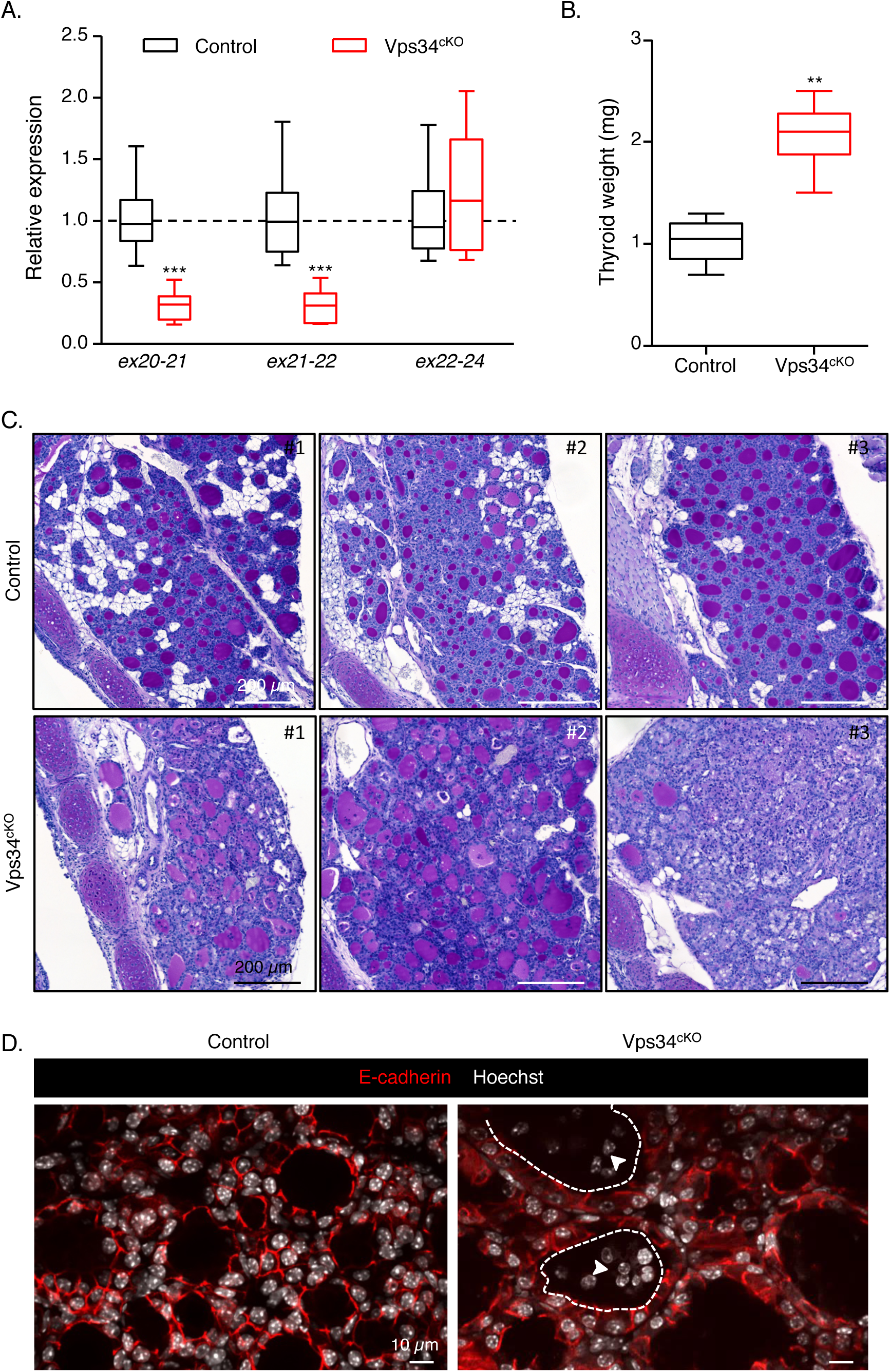
Genetic and histopathological characterization. ***A. Extensive genetic excision of Vps34 exon 21***. Compared with control (black boxes), Vps34^cKO^ thyroid (red boxes) shows ∼70% reduction in exon 21 mRNA level at P14. The unchanged mRNA level spanning exons 22-24 serves as control. Boxes with median and percentiles of 10 WT and 11 cKO samples; ***, p< 0.0001 by Mann-Whitney non-parametric test. ***B. Increased thyroid weight.*** Compared with control (black boxes), Vps34^cKO^ thyroid (red boxes) shows a two-fold increase in thyroid weight. Boxes with median and percentiles of 8 WT and 6 cKO samples; **, p< 0.01 by Mann-Whitney non-parametric test. ***C. Histopathological evidence for colloid exhaustion*.** As compared to control thyroid tissues (n=3) where all follicles show regular lumen filling with homogenous and intense PAS (Periodic Acid Shiff) staining, Vps34^cKO^ thyroids (n=3) present fewer, mostly centrally located, PAS-stained follicles and with weaker staining intensity, other follicles that appear empty (for further quantification, see Supplementary Figure 1). ***D. Vps34^cKO^ thyrocytes are altered and follicles contains abundant non-epithelial cells***. Nuclei are labeled by Hoechst (shown in white); thyrocyte basolateral contours are labeled for E-cadherin (red); two lumen boundaries are delineated by broken lines. As compared to control thyroid follicles, Vps34^cKO^ follicular structures are frequently irregular and containing additional cells inside the lumen (arrowheads). These cells are not labeled for E-cadherin.

### Vps34^cKO^ thyroids show signs of hyperstimulation by thyroid stimulating hormone (TSH)

At P14, thyroid glands from Vps34^cKO^ were almost twice heavier as compared to control glands (Fig. 1B). Histological staining using Periodic-Acid-Schiff (PAS, which in the thyroid reflects production of the thyroglobulin glycoprotein) readily revealed striking differences between thyroid glands of control and Vps34^cKO^ mice (Fig. 1C). Thyroid glands of control mice appeared as assemblies of round follicular sections of variable diameter, filled with colloid intensely and homogenously stained by PAS. Although histology of Vps34^cKO^ was more variable, the three representative individual thyroids shown in Fig. 1C revealed irregular follicle shape, especially at the periphery of the gland, and weaker or absent PAS staining (Fig. 1C and Suppl. Fig. 1). In addition, the PAS-negative follicular space were often filled with nuclei (see Vps34^cKO^ #3). Quantification revealed that 90% of control follicles had a regular shape and intense (+++) PAS staining (Suppl. Fig. 1). On the contrary, only 50% of Vps34^cKO^ follicles displayed a regular shape and most of the follicles were weakly positive for PAS (++ or +) (Suppl. Fig. 1). In addition, 10 to 50% of the Vps34^cKO^ follicles showed no PAS staining (-) and luminal cells were found in 40-70% of the follicles (Suppl. Fig. 1). Confocal immunofluorescence microscopy for the epithelial marker E-cadherin confirmed heterogeneity of Vps34^cKO^ follicles, but also of thyrocytes. Hoechst labelling revealed the presence of several nuclei in a significant fraction of lumina (Fig. 1D). These histological features of follicle remodelling and colloid consumption up to exhaustion, suggested perturbed thyroid function.

### Vps34^cKO^ causes severe non-compensated hypothyroidism

To directly test the hypothesis that Vps34^cKO^ could impair thyroid hormone production, plasma was collected at P14 and analyzed for the levels of T_4_ and TSH. In P14 control pups, values were 6.4 ± 1.5 μg/dL for T_4_, and 75 ± 110 mU/L for TSH (Fig. 2). In Vps34^cKO^ mice, we found extremely low T_4_ values (0.75 ± 0.62 μg/dL) and very high plasma levels of TSH (19,311 ± 10,482 mU/L) (Fig. 2). These data indicate severe, non-compensated hypothyroidism. This observation, compatible with the growth retardation observed after P15 (19), could be explained by mislocalization of one or several basolateral and/or apical actors involved in thyroid hormone synthesis, as we reported in Vps34^cKO^ kidney proximal tubular cells (19).

**Figure 2.**
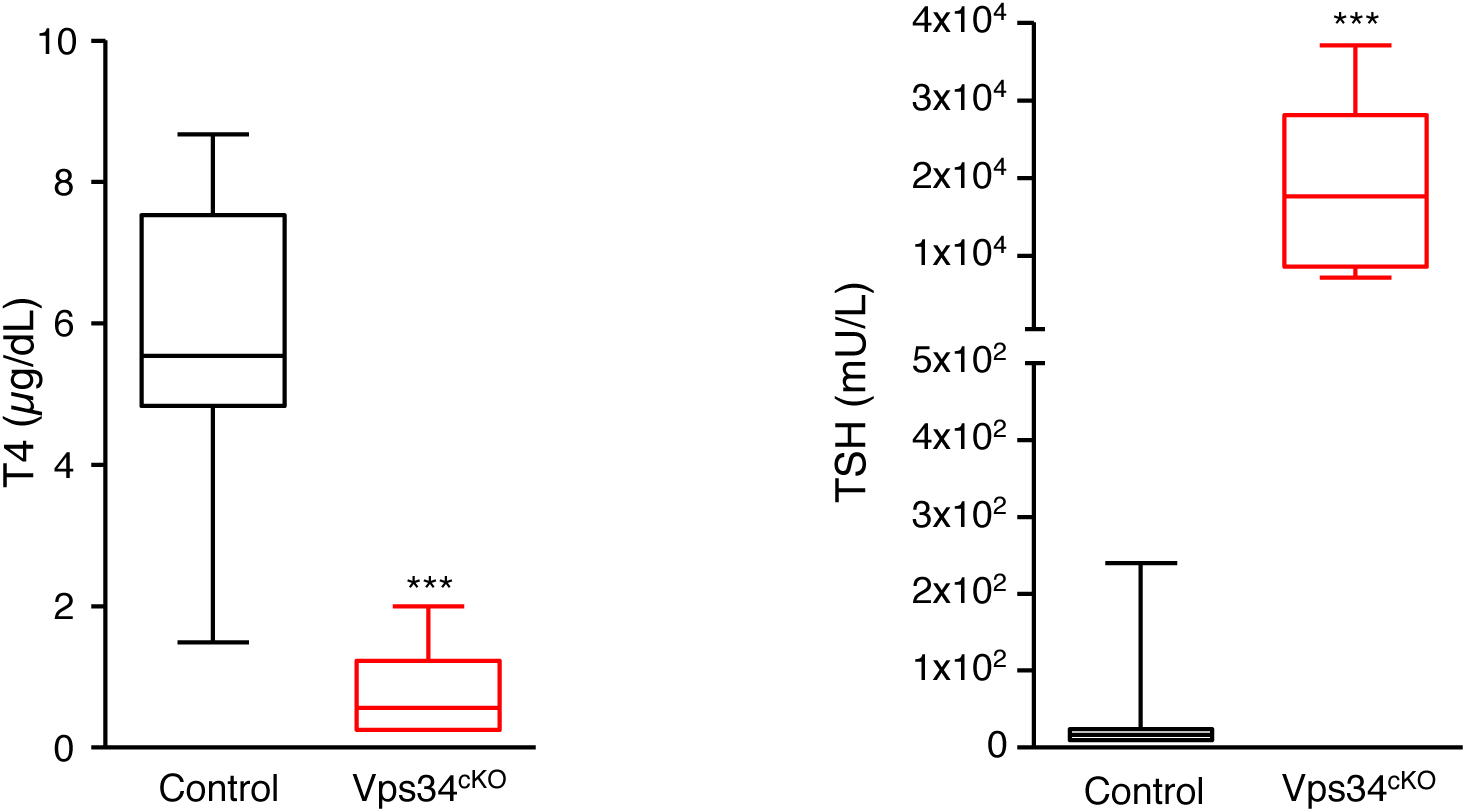
Vps34^cKO^ mice display severe hypothyroidism associated with high TSH levels. As compared to control mice (black boxes), T_4_ plasma level in Vps34^cKO^ is extremely low (red boxes). Conversely, TSH levels are dramatically elevated. (10 WT and 9 cKO samples; ***, p= 0.0003 by Mann-Whitney non-parametric test).

### Vps34 cKO display reduced iodine organification

The reduced or absent PAS staining (Fig. 1C) and the low T_4_ plasma levels in Vps34^cKO^ mice (Fig. 2) might be caused by defective basolateral iodine uptake, apical transport and/or apical organification on thyroglobulin. Due to lack or very poor specificity of antibodies directed against murine proteins involved in these processes, we first measured their mRNA expression levels in total thyroids at P14 (Fig. 3). We observed no change in mRNA expression of *Nis*, *Ano1*, *Tg*, *Tpo* and *Duox2,* a significant increase in *Tshr* and *Slc26a4* expression, and a significant decrease of *Slc26a7* and *Duoxa2* mRNA levels. At this stage, we cannot conclude if these changes contribute to the observed hypothyroidism of Vps34^cKO^.

**Figure 3.**
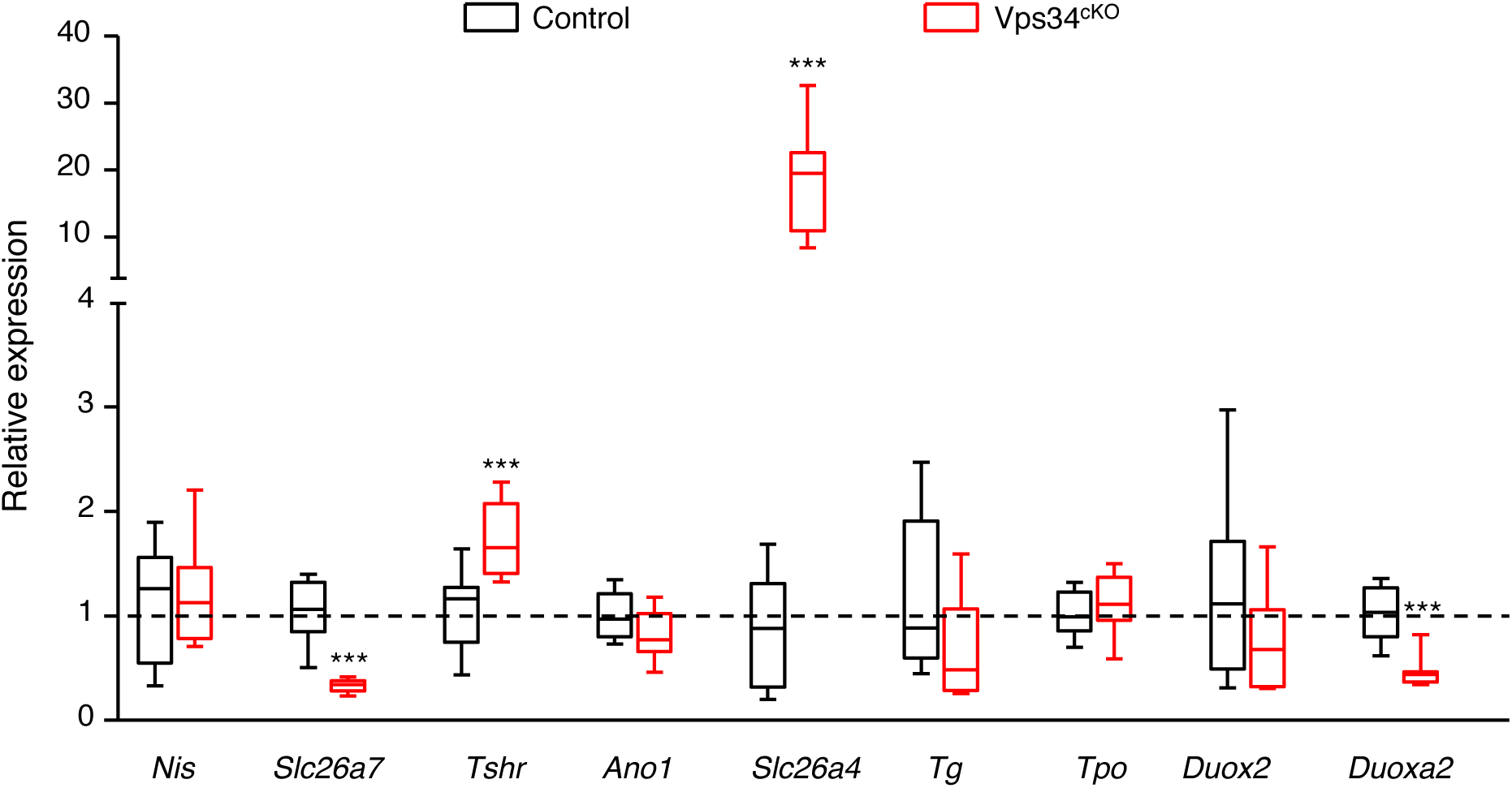
Relative expression level of main actors of thyroid hormonogenesis at P14. Gene expression analysis by RT-qPCR, presented as boxes with median and percentiles. As compared to control thyroid (black boxes), expression in Vps34^cKO^ (red boxes) is preserved for the thyroid-specific genes *Nis*, *Tpo*, *Tg* and *Duox2.* Expression of the TSH receptor (*Tshr*) and of *Slc26a4* is significantly increased in Vps34^cKO^, while expression of *Slc26a7* and *Duoxa2* is significantly decreased. Boxes with median and percentiles of at least 9 control and 9 Vps34^cKO^ samples; ***, p< 0.001 by Mann-Whitney non-parametric test.

To further test whether basolateral NIS and SLC26A7 and apical ANO1, SLC26A4, TPO, DUOX and DUOXA were all correctly localized, we functionally assayed their combined activity by injecting iodine-deprived pups at P9 with ^125^I. At P10, 24-h post-injection, thyroid lobes were collected and radioactivity measured before and after protein precipitation by TCA (Fig. 4A). ^125^I uptake was not statistically-different in Vps34^cKO^ and the three control genotypes (Fig. 4A), suggesting normal NIS and SLC26A7 function and thus basolateral localization. In marked contrast, only 10% of thyroid ^125^I was bound to proteins in Vps34^cKO^ compared to approximately 55% in control genotypes at this stage (Fig. 4A). This indicates a major effect of Vps34 on one or several apical proteins involved in iodine organification.

**Figure 4.**
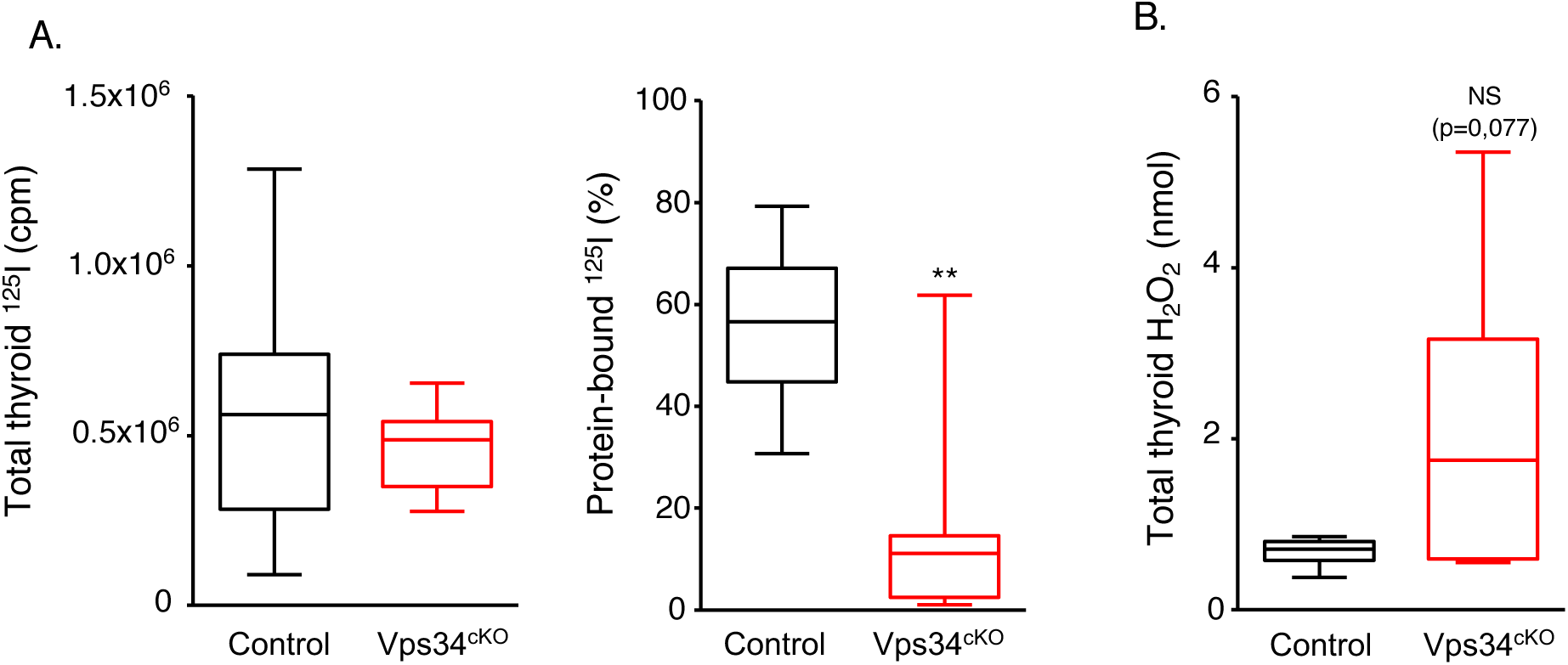
Normal uptake of iodine but major defect of organification in Vps34^cKO^ thyroid. ***A.*** Comparable ^125^I-iodine uptake in the thyroid in control genotypes (black box) and Vps34^cKO^ thyroid (red box), but much reduced ^125^I bound to protein (in %) in Vps34^cKO^ thyroid, suggesting defective organification. Boxes with median and percentiles of 39 (10 Flox/+, 19 Flox/Flox, 10 Cre;Flox/+) and 12 Vps34^cKO^ (Cre;Flox/Flox) samples; ***, p< 0.001 by Mann-Whitney non-parametric test. **B**. Level of H_2_O_2_ in total thyroid extracts show a trend to an increase in Vps34^cKO^ thyroid. Boxes with median and percentiles of 9 control and 9 Vps34^cKO^ samples; p= 0.077 by Mann-Whitney non-parametric test.

Normal *Ano1* and increased *Slc26a4/Pendrin* mRNA levels suggested correct, or even improved, transfer of iodine in the follicular lumen. On the other hand, decreased *Duoxa2* might impact on H_2_O_2_ production. We thus measured the production of H_2_O_2_ in control and Vps34^cKO^ thyroid lobes. Surprisingly, the median level of H_2_O_2_ production in Vps34^cKO^ was higher, even if we observed variability (Fig. 4B), indicating that the DUOX/DUOXA pair is functional in Vps34^cKO^ thyroid. Altogether, these results suggest defective localization of one or several apical actors involved in iodine organification.

### Apical polarity is impaired in Vps34^cKO^

Apico-basal polarity is essential for thyrocyte function and defects in polarity might impact on the delivery, and thus localization, of actors involved in thyroid hormone synthesis. As general markers to assess polarity, we used the basement membrane protein laminin, the basolateral Na^+^/K^+^-ATPase, E-cadherin and 11-catenin, the apically-localized ERM family member ezrin and the tight junction-associated protein ZO-1. Control thyrocytes displayed a well-defined apico-basal polarity with laminin assembled on the basal side, with Na^+^/K^+^-ATPase, E-cadherin and □-catenin restricted to the basolateral membrane and separated from apical ezrin by the tight junction, visualized by ZO-1 (Fig. 5A and 5B). In Vps34^cKO^ thyrocytes, basal laminin was correctly assembled and Na^+^/K^+^-ATPase, E-cadherin and □-catenin were restricted to the basolateral membrane, indicating the presence of functional tight junctions. However, on the apical side of Vps34^cKO^ thyrocytes, the ezrin signal was weaker or absent from the lumen-facing pole of some thyrocytes (Fig. 5A), and most cells also lacked ZO-1 protein labelling (Fig. 5B). These data suggest impaired apical polarity in Vps34^cKO^, which might contribute to defective localization of apical actors involved in iodine organification.

**Figure 5.**
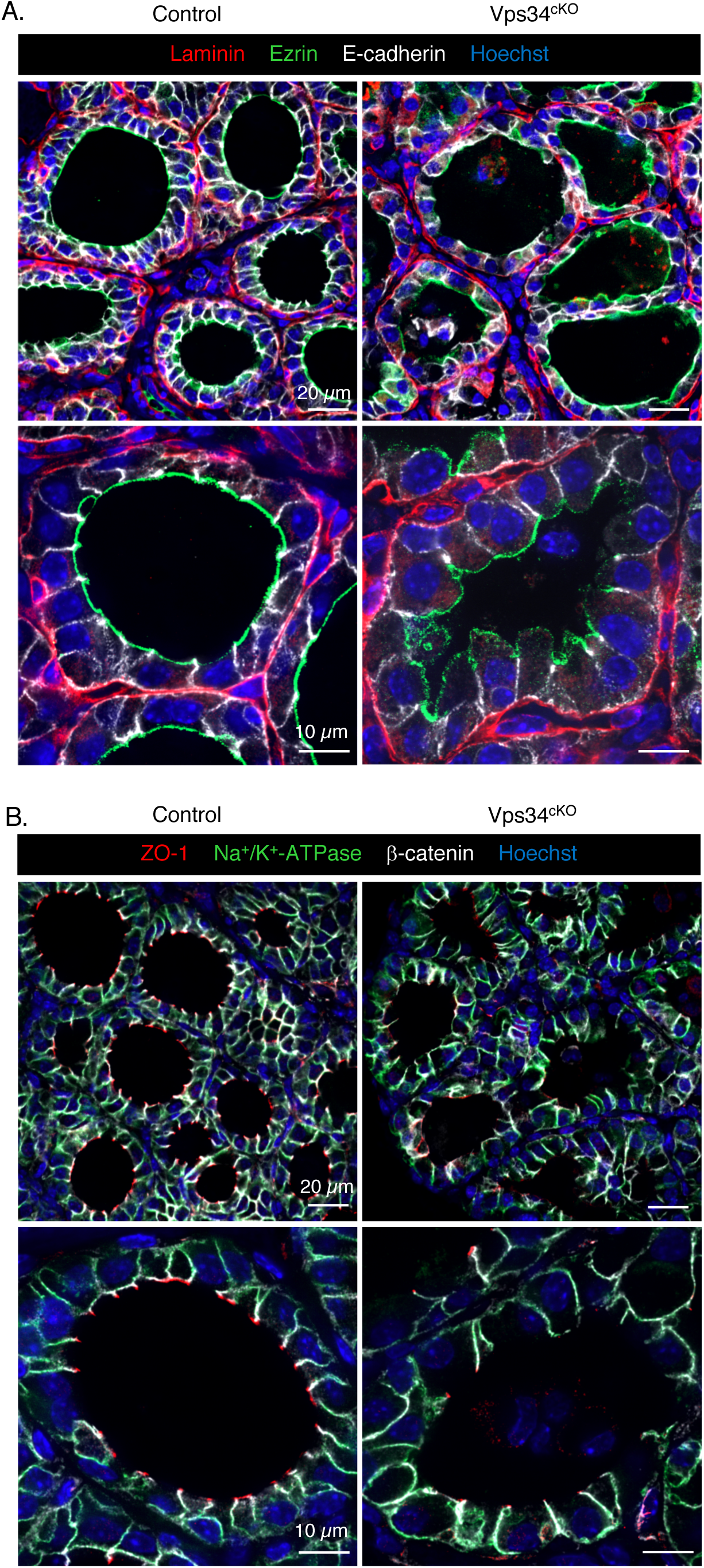
Normal basolateral but impaired apical organization of tyrocytes. ***A.*** Thyroid sections from control (left) and Vps34^cKO^ (right) labeled for laminin (red), ezrin (green) and E-cadherin (white). Nuclei are labeled by Hoechst (shown in blue). In control thyroid, laminin surrounds follicles composed of thyrocytes delineated by basolateral E-cadherin and apical ezrin. In Vps34^cKO^, laminin normally surrounds follicles composed of thyrocytes with well-defined basolateral E-cadherin but weaker and less regular apical ezrin. Note here apical membrane bulging in the colloidal space. ***B.*** Thyroid sections from control (left) and Vps34^cKO^ (right) labeled for ZO-1 (red), Na^+^/K^+^-ATPase (green) or β-catenin (white). Nuclei are labeled by Hoechst (shown in blue). In control and Vps34^cKO^ thyroid, Na^+^/K^+^-ATPase and β-catenin are correctly localized and restricted to the basolateral pole of the thyrocytes, indicating the presence of a tight junction. However, the tight junction-associated protein ZO-1 is only detected at few apico/basolateral junctions.

### Vps34^cKO^ mice display defective thyroglobulin iodination

To further analyse the significance of the lack of PAS staining in a fraction of Vps34^cKO^ follicles (Fig. 1C) and the defect in iodine organification (Fig. 4), we assessed thyroglobulin protein expression and its associated T_4_ hormonogenic peptide (I-Tg) by immunofluorescence. Low magnification of control sections showed an identical distribution pattern of thyroglobulin and I-Tg, homogeneously filling all round colloidal spaces (Fig. 6A). In Vps34^cKO^, the thyroglobulin labelling was more heterogeneous, mainly due to the presence of cells in the colloid. Remarkably, antibodies recognizing the T_4_ hormonogenic peptide often failed to label colloidal spaces even when containing Tg in the adjacent section (Fig. 6A), thereby confirming the organification defect (Fig. 4A). Quantification revealed that only a quarter (25.2 ± 11.9 %) of the follicles were labeled for T_4_ hormonogenic peptide. Of specific interest, whereas the Tg signal was restricted to the follicle lumen in control thyroid, high magnification revealed that the Tg signal was frequently seen within Vps34^cKO^ thyrocytes (Fig. 6B, arrowheads). These observations indicated that Vps34 deletion caused defective thyroglobulin iodination, as well as reduced Tg exocytosis and/or excessive Tg endocytosis.

**Figure 6.**
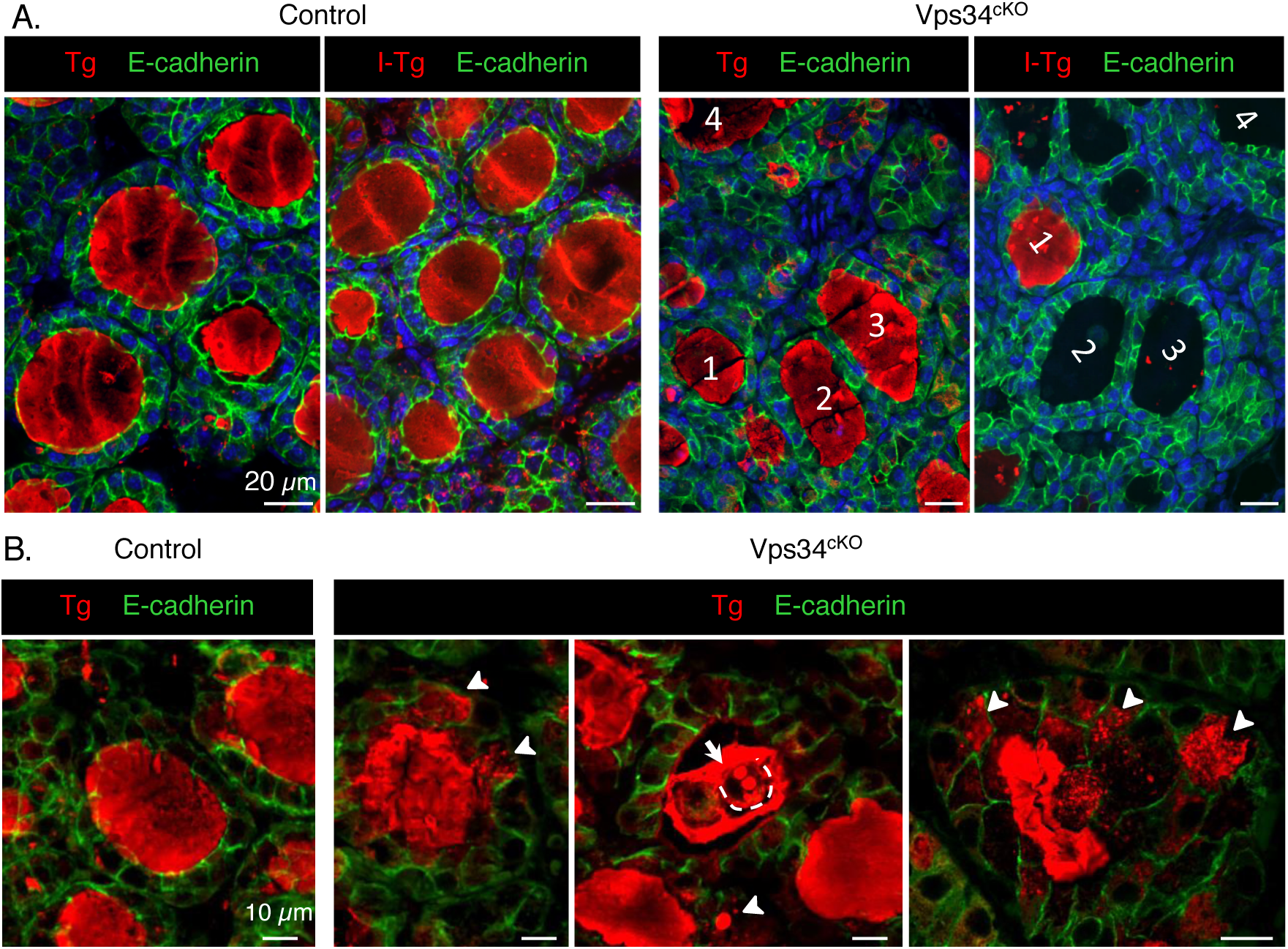
Evidence for defective iodide organification in Vps34^cKO^ thyroid: follicular lumina with Tg but devoid of iodinated-Tg. **A.** Thyroid sections from control (left) and Vps34^cKO^ (right) labeled for E-cadherin (green) and thyroglobulin (red), recognized either for a core protein epitope (Tg), or a hormonogenic peptide (I-Tg). Control follicular lumina are round and uniformly labeled for both Tg and I-Tg. In Vps34^cKO^, follicular structures are less regular, with the majority containing Tg, but not the hormonogenic peptide. In Vps34^cKO^, two serial (tilted) sections are shown. ***B***. ***Comparison of Tg labelling in control and Vps34******^cKO^ thyrocytes.*** In control follicles, Tg is essentially restricted to lumen, with little or no signal inside thyrocytes. In Vps34^cKO^, Tg is readily detected within thyrocytes as collections of submicrometric dots or larger spheres (arrowheads). Of note, some luminal cells (broken line in middle panel) also contain Tg-labeled spheres, indicating active endocytosis (arrow).

### Impaired lysosomal function and I-Tg proteolysis in Vps34^cKO^ thyroid

Vps34 is involved in endocytic trafficking and cKO of Vps34 in postnatal kidney glomeruli podocytes causes a strong increase of the overall immunofluorescence signal for the lysosomal membrane marker LAMP-1, indicating enhanced lysosome biogenesis (29, 30). We further reported that absence of Vps34 in kidney PTCs causes an increase in the actual size of lysosomes, that were sometimes enlarged, and mislocalized toward the basal pole of the cell (19). Given that lysosomal proteases in thyrocytes are important to excise hormonogenic peptides to release free T_3_ and T_4_, we investigated LAMP-1-labeled structures in Vps34^cKO^ thyrocytes as a proxy for lysosomal function, and its co-localization with Tg and I-Tg to demonstrate endocytosis. In our conditions, immunofluorescence on control thyroid sections only produced a weak LAMP-1 signal in the E-cadherin-positive epithelial thyrocytes (Fig. 7A and 7B). On the contrary, a strong and widespread LAMP-1 signal was observed in all Vps34^cKO^ thyrocytes. As shown in the enlarged boxes of Fig. 7A and 7B (separate emission channels in white), LAMP-1-positive late endosomes/lysosomes were more abundant, and sometimes enlarged in Vps34^cKO^ thyrocytes as compared to control. In addition, the intracellular Tg protein, found in most Vps34^cKO^ thyrocytes (Fig. 7A, arrowheads), most often colocalized with LAMP-1-positive structures. This indicates that colloidal Tg was endocytosed by thyrocytes, but that trafficking to or Tg proteolysis within the lysosomes was slower or impaired, as compared to controls. Of note, luminal cells also contained LAMP-1 structures positive for Tg (Fig. 7A, right).

**Figure 7.**
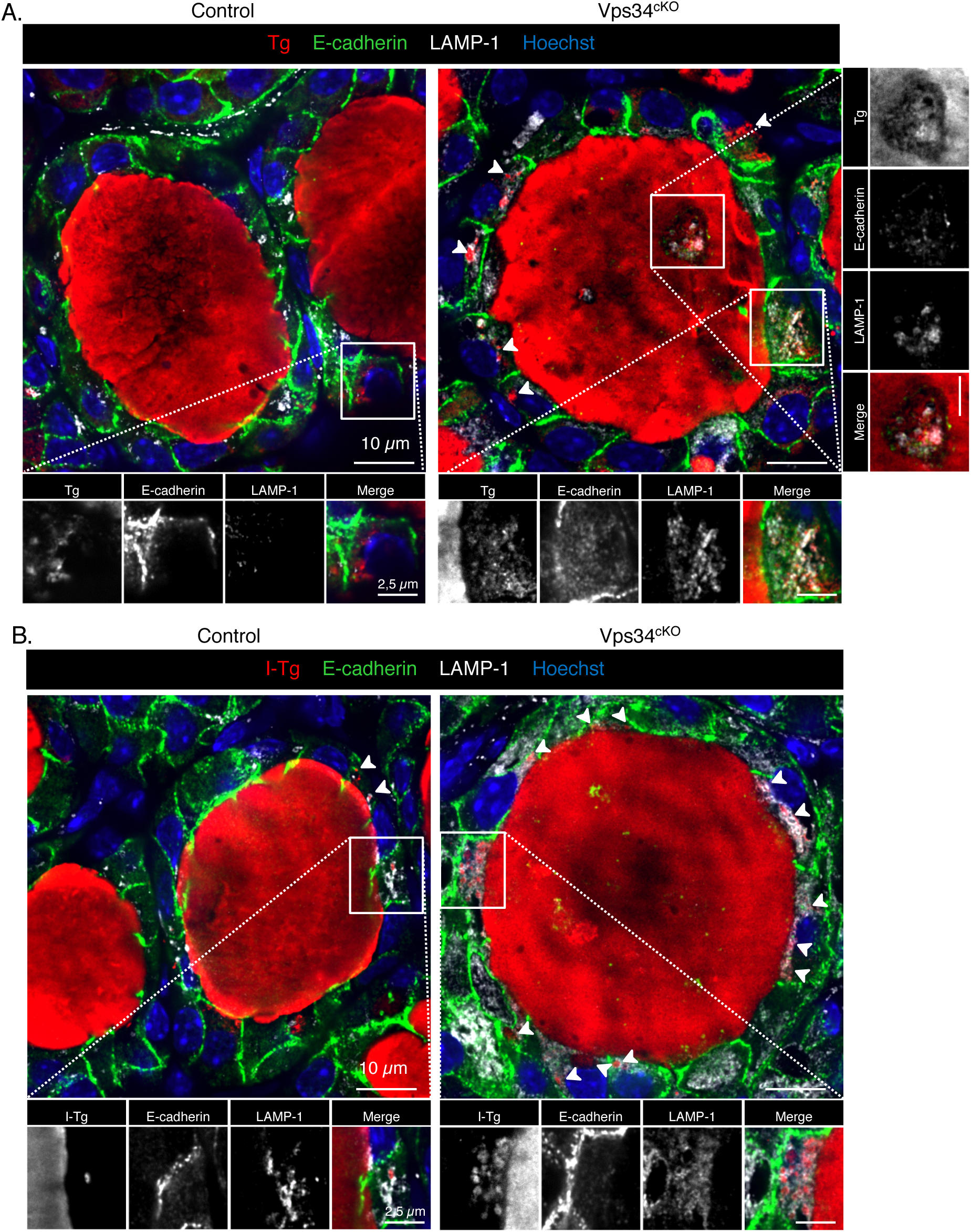
Vps34^cKO^ thyrocytes display increased LAMP-1 signal and impaired I-Tg proteolytic processing. **A.** Thyroid sections from control (left) and Vps34^cKO^ (right) labeled for thyroglobulin (red), E-cadherin (green) and lysosomal LAMP-1 (white). Nuclei are labeled by Hoechst (shown in blue). In control, thyrocytes rarely contain intracellular Tg and LAMP-1 signal is weak. In Vps34^cKO^, most thyrocytes have intracellular Tg (arrowheads) and LAMP-1 signal is much increased. Insets show magnification with separate emissions, then merged, channels. Note colocalization of Tg with LAMP-1 in luminal cells (at right). **B.** Thyroid sections from control (left) and Vps34^cKO^ (right) labeled for iodinated-thyroglobulin (I-Tg, red), E-cadherin (green) and lysosomal LAMP-1 (white). Nuclei are labeled by Hoechst (shown in blue). As compared to control follicles which show a weak signal for LAMP-1, and rare intracellular I-Tg, LAMP-1 signal is intense in Vps34^cKO^ thyrocytes and intracellular I-Tg dots are frequently observed (arrowheads). As shown in the enlargements below, LAMP-1-labeled structures in Vps34^cKO^ thyrocytes are enlarged or vacuolated as compared to control.

Additional supporting evidence for impaired lysosomal proteolysis came from the analysis of I-Tg (Fig. 7B). Indeed, Vps34^cKO^ thyrocytes often contained intracellular signal for I-Tg, colocalizing within LAMP-1 structures (Fig. 7B, enlarged boxes,). Thus, in addition to reduced iodine organification, defective lysosomal proteolysis of T_3_/T_4_, if/when hormonogenic peptides were still formed on Tg, was also accounting for the low T_3_/T_4_ plasma levels observed in Vps34^cKO^ (Fig. 2).

### Evaluation of autophagy in Vps34^cKO^ thyroid

The deletion of Vps34 in the kidney also causes a block of autophagy (19). We therefore assessed the expression of p62 (also called sequestosome-1, or SQSTM1), a polyubiquitin-binding protein that interacts with LC3b (microtubule associated protein 1 light chain) on the autophagosome membrane and is normally continuously degraded by the autophagy process (31). Since p62 accumulates when completion of autophagy is inhibited, p62 can be used as a marker to study autophagic flux (32). Although we found a weak punctiform LC3 signal in control thyrocytes, we observed no p62 signal, indicating normal autophagic flux in control thyroid. As reported, few LAMP-1-labelled structures were found in control thyroid. In contrast, much larger structures, mostly double-labeled for p62 and LC3, were easily detected in the cytoplasm of Vps34^cKO^ thyrocytes. As the LAMP-1 signal was increased in Vps34^cKO^, these p62/LC3-positive punctae sometimes co-localized with LAMP-1. We concluded that LC3 could be recruited on p62 aggregates but that progression to autophagosome maturation and fusion with, and degradation by, lysosomes was arrested. This suggested that deletion of Vps34 in thyrocytes abrogated the autophagic flux.

Recent evidences have demonstrated that p62 is also an activator of the antioxidant KEAP1/NRF2 pathway, which impacts on thyrocyte physiology and Tg biology (33). We thus examined the possibility that stabilized p62 would compete with NRF2 for KEAP1 binding in Vps34^cKO^, thus allowing NRF2 to reach the nucleus and activate gene expression. We quantified the mRNA levels of *Nrf2* and its target genes *Nqo1*, *Gpx2* and *Txnrd1*. Although we measured a slight decrease of *Nrf2* mRNA in Vps34^cKO^, two of its target genes, namely the quinone reductase, *Nqo1*, and the thioredoxin reductase 1, *Txnrd1*, were upregulated 1.9- and 3.5-fold, respectively (Fig. 8B). This observation supports the hypothesis that KEAP1 interaction with p62 is favored in Vps34^cKO^ and that NRF2 is thus displaced and able to migrate to the nucleus to regulate gene expression.

**Figure 8.**
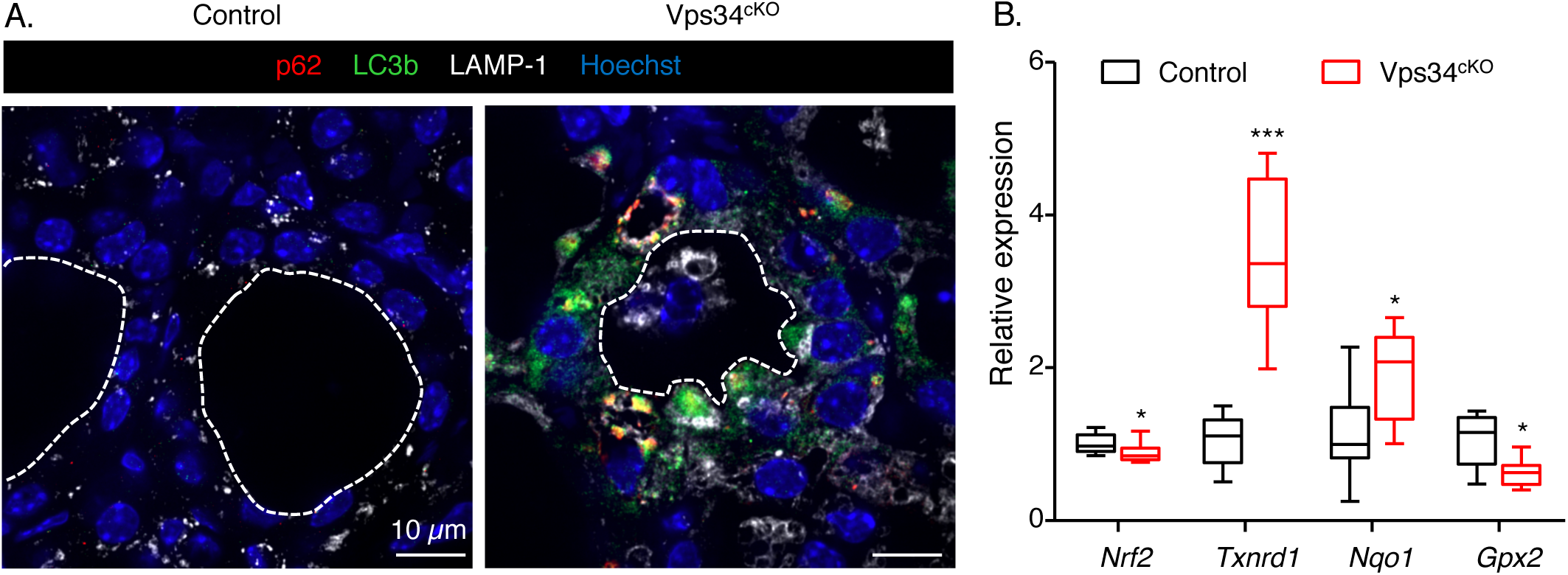
Vps34^cKO^ thyrocytes show strong accumulation of p62 and LC3b dots. **A.** Thyroid sections from control (left) and Vps34^cKO^ (right) labeled for p62 (red), LC3b (green) and lysosomal LAMP-1 (white). Nuclei are labeled by Hoechst (shown in blue). Compared with control follicles, which show low LAMP-1 and almost no signal for p62 and LC3b, all three markers are very strong in Vps34^cKO^ thyroid sections. p62 perfectly co-localizes with LC3b and labels large aggregates. These aggregates are adjacent to LAMP-1-positive structures. **B. Increased NRF2 signaling.** Gene expression analysis by RT-qPCR, presented as boxes with median and percentiles. As compared to control thyroid (black boxes), expression of *Nrf2* is reduced in Vps34^cKO^ (red boxes), but level of two out of three NRF2 target genes, *Txnrd1* and *Nqo1* is increased. Boxes with median and percentiles of at least 9 control and 9 Vps34^cKO^ samples; *, p< 0.05 and ***, p< 0,001 by Mann-Whitney non-parametric test.

We also evaluated macroautophagy/mitophagy by immunolocalizing TOM20 and LAMP-1. In Vps34^cKO^ thyrocytes, TOM20-labeled structures did not colocalize with the more abundant, enlarged LAMP-1+ structures (Suppl. Fig. 2A), as we also reported in Vps34^cKO^ kidney proximal tubular cells (19). This suggested that sequestration of mitochondria into autophagosomes is either not triggered, which is unlikely for altered cells, or abortive at P14 in Vps34^cKO^ thyrocytes, and/or that fusion of autophagosomes with lysosomes is prevented. We then evaluated the chaperone-mediated autophagy by immunolocalizing LAMP-2A. As compared to control thyroid, LAMP-2A-labeled structures were more abundant in Vps34^cKO^ thyrocytes (Suppl. Fig. 2B), as we also reported in Vps34^cKO^ kidney proximal tubular cells (19).

### Luminal cells present in Vps34^cKO^ follicles are macrophages

Finally, we investigated the origin of the cells present in the colloidal space. As observed in Figure 7A, cells trapped in the follicular lumen showed a strong LAMP-1 signal (Fig. 7A). However, they were negative for E-cadherin (Figs. 1D and 5A). In addition, we found that luminal cells were also negative for the permanently-expressed thyrocyte-specific transcription factor, TTF-1 (Fig. 9A), indicating that these cells were not derived from the thyrocyte lineage. Lineage-tracing experiments on Pax8-Cre; Vps34^fl/fl^;ROSA-STOP-YFP pups at P14 further revealed YFP-positive signal only in follicle-delineating thyrocytes and not in the luminal cells (Fig. 9B). Thus, luminal cells had never expressed Pax8 and were thus not derived from thyrocyte progenitors. Although infiltration by C-cells was a possibilty, we failed to label luminal cells for the Prox1 transcription factor. As luminal cells displayed LAMP-1 signal and were also positive for Tg (Fig. 7A), we tested the possibility that luminal cells were infiltrating macrophages. This was the case: luminal cells were labeled by the macrophage marker F4/80 (Fig. 10A), but not by E-cadherin. To confirm that colloid was consumed by infiltrated macrophages, we performed a triple immunolabelling for F4/80, LAMP-1 and I-Tg. Luminal F4/80-positive cells displayed huge LAMP-1-positive lysosomes filled with I-Tg (Fig. 10B). Thus, colloid consumption (Tg in Fig. 7A and I-Tg in Fig. 9D) by infiltrated macrophages provides a third explanation for the low T_4_ plasma level observed in Vps34^cKO^.

**Figure 9.**
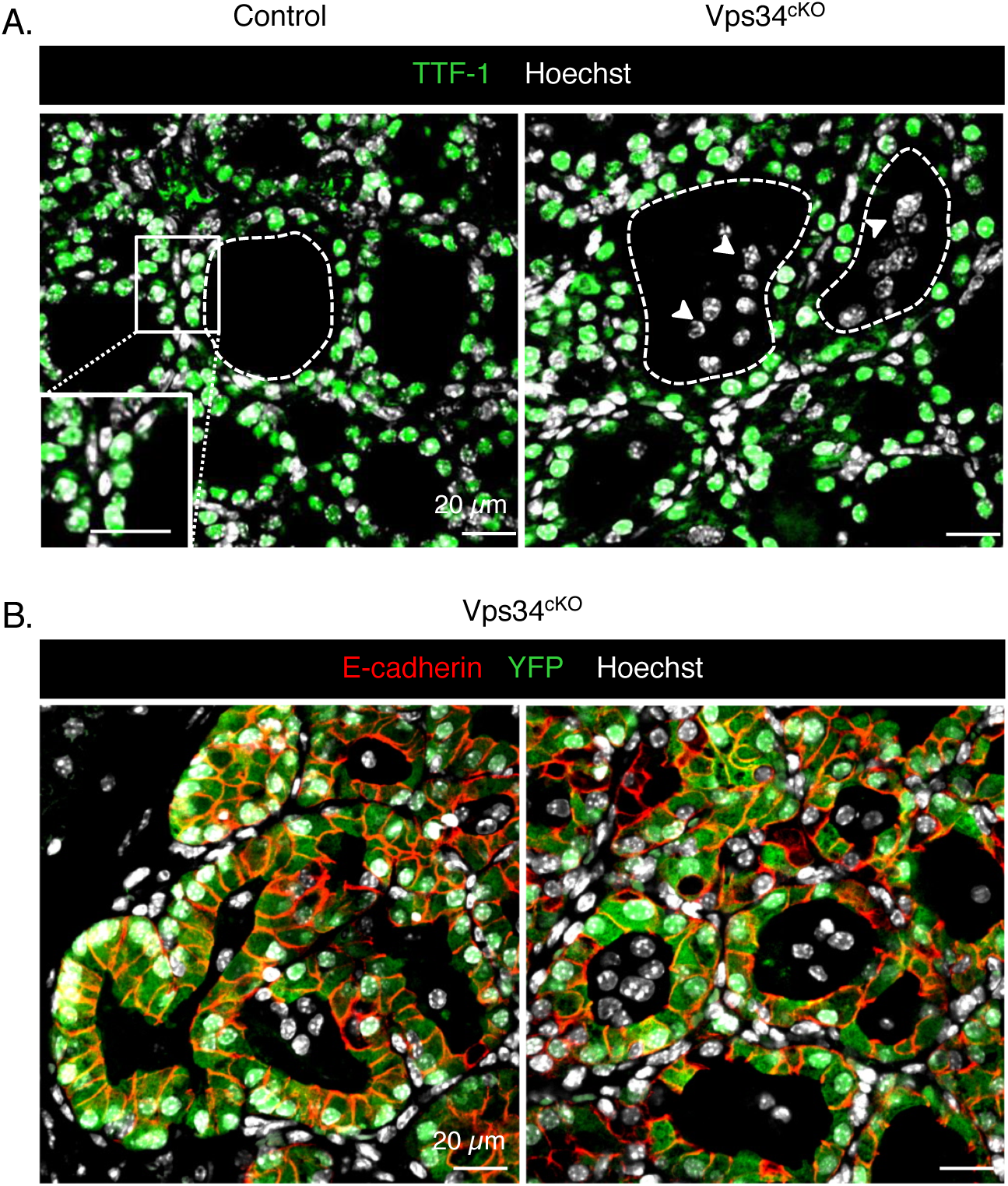
Luminal cells in Vps34^cKO^ thyroid are not thyrocytes. ***A.*** Immunolabelling for thyrocyte transcription factor-1 (TTF-1, green); nuclei are visualized with the Hoechst stain (shown in white). In both control and Vps34^cKO^ thyroid, nuclei of thyrocytes circumscribing the follicular lumina are all labeled by the thyrocyte-specific transcription-factor-1, TTF-1. As indicated by the arrowheads, nuclei of luminal cells show no signal for TTF-1 (only white signal representing Hoechst). ***B.*** Immunolabelling for E-cadherin (red) and YFP (green); nuclei are visualized with the Hoechst stain (shown in blue). Representative images of Vps34cKO thyroid sections reveal expression of YFP only in cells surrounding the follicular lumina, i.e. in cells where Pax8-Cre has been active. Luminal cells do not derive from thyrocyte progenitors as they are negative for YFP.

**Figure 10.**
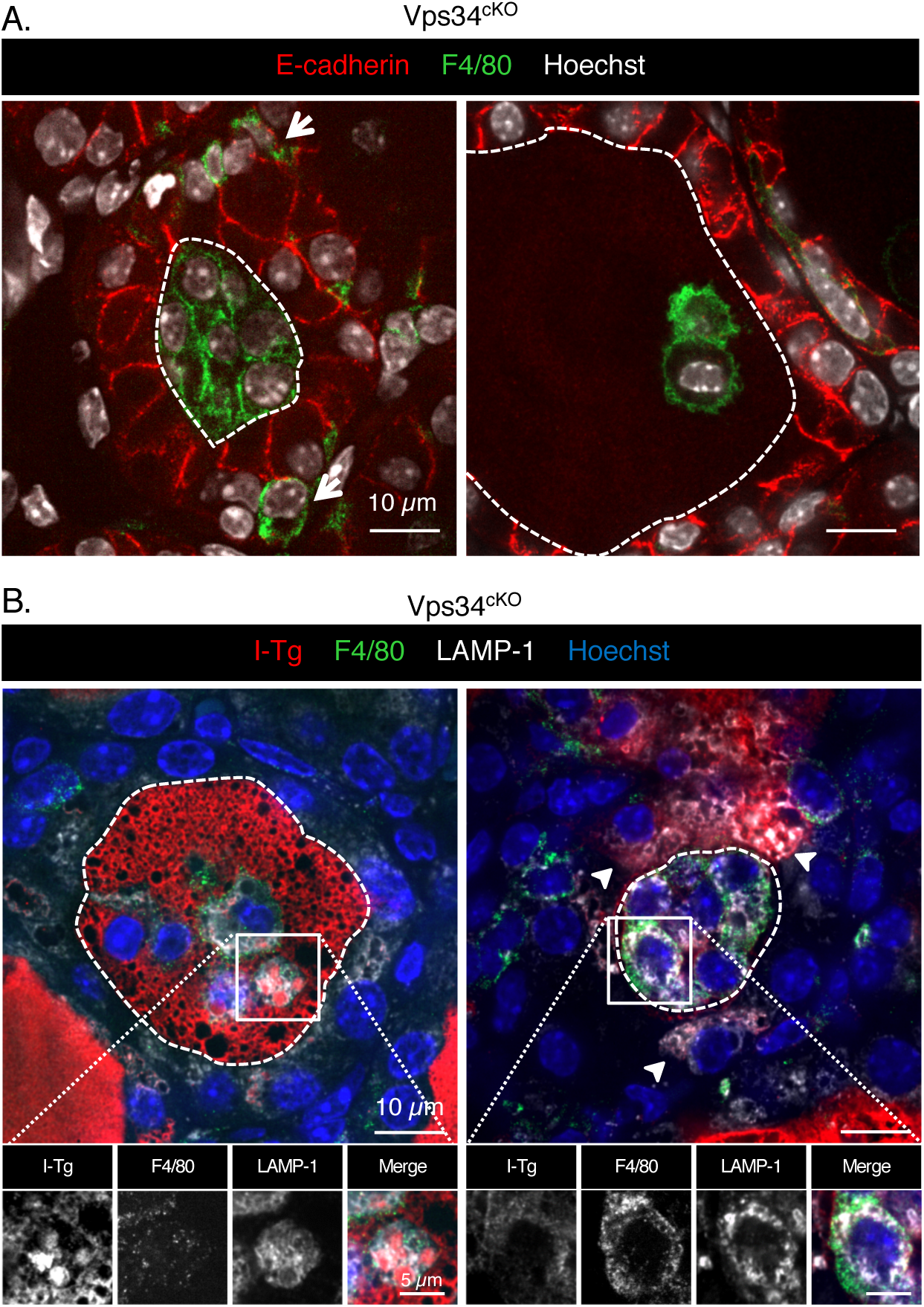
Luminal cells in Vps34^cKO^ thyroid are macrophages. ***A.*** Immunolabelling for E-cadherin (red), and for the conventional macrophage marker (F4/80, green); nuclei are visualized with the Hoechst stain (shown in white). In Vps34^cKO^ thyroid, F4/80 signal can be found in the interstitium (arrows), but also in the colloidal space, thus identifying luminal cells as infiltrating macrophages. **D. Luminal cells are taking up the colloid.** Immunolabelling for I-Tg (red), F4/80 (green) and LAMP-1 (white). Nuclei are visualized with the Hoechst stain (shown in blue). Luminal cells in Vps34^cKO^ have abundant LAMP-1 structures that surround I-Tg droplets. Arrowheads show additional I-Tg in thyrocytes. Broken lines indicate luminal contours.

## Discussion

In this study, we report that deleting Vps34 in thyrocytes by Pax8-Cre-driven recombination causes several defects in the thyroid: (i) doubling of thyroid weight and perturbed thyroid parenchyma organization, with reduced PAS^+^ colloidal spaces; (ii) severe hypothyroidism with collapsed plasma T_4_ levels and very high TSH; (iii) strong decrease of ^125^I organification, at comparable ^125^I uptake, and of T_4_ formation on thyroglobulin (detected by immunofluorescence as early as P3); (iv) defective apical polarization; (v) impaired lysosomal proteolysis; (vi) infiltration of macrophages in the colloid. Some of these features, combined with delayed “eye opening” and impaired postnatal growth after two weeks (18), phenocopy the impact of *Duoxa2* KO in the thyroid (34).

DUOXA2 is a chaperone protein required for the correct localization of DUOX2 at the apical pole of thyrocyte, where the complex (DUOX2/DUOXA2) produces H_2_O_2_. In the absence of DUOXA2, H_2_O_2_ is not produced and subsequent thyroperoxidase-mediated oxidation of iodide into reactive compounds and thyroglobulin iodination are abolished (34). The similarity of the severity of hypothyroidism in Vps34^cKO^ and *Duoxa2* KO mice prompted us pay a closer look at DUOXA2 in Vps34^cKO^. We found that the expression of *Duoxa2* started decreasing after P3, reaching two-fold lower values at P14. It is interesting to note that expression of *Duox2* and *Duoxa2* mRNAs was quantitatively different, despite of the fact that they share the same promoter (35). Whether decreased *Duoxa2* mRNA expression is reflected by equivalent two-fold decrease of DUOX2/DUOXA2 complex at the apical pole is unknown. However, it is very unlikely that a two-fold decrease would by itself so severely impact on thyroglobulin iodination, since heterozygous mice for *Duoxa2* deletion have virtually no phenotype. In addition, our measurements of H_2_O_2_ levels in total Vps34^cKO^ thyroid extracts also support normal, or increased, function of the DUOX2/DUOXA2 complex. Instead, based on the known functions of Vps34 in kidney PTCs, where its inactivation causes lack of apical localization of endocytic receptors (megalin, cubilin) and solute transporters (NaPi-IIa, SGLT-2) (19), it would be extremely interesting to localize actors of thyroid hormonogenesis on thyroid sections, and assess their basolateral or apical addressing in Vps34^cKO^ thyrocytes. Unfortunately, reliable antibodies to detect most of these proteins are not yet available for mice. Nevertheless, ^125^I uptake and processing experiments suggests normal localization and function of the sodium-iodine symporter, NIS, and/or the alternative transporter SLC26A7 (3), and rather support the hypothesis of defective apical localization of one or several actors involved in thyroid hormonogenesis (Ano1, pendrin, TPO, DUOX, DUOXA).

Characteristic histopathological alterations combined with the very high plasma TSH levels and two-fold increased *Tshr* expression in Vps34^cKO^ thyroid suggest that thyrocytes are in a hyperstimulated state. For example, DuoxA2 KO, which display very high TSH levels, present a 20- and 5-fold higher expression level of *Nis* and *Tpo*, as compared to controls (34). However, the expression of these two sensitive target genes of the TSH signaling pathway, *Nis* and *Tpo*, was surprisingly unchanged in Vps34^cKO^, arguing against thyrocyte hyperstimulation. We favor the hypothesis of TSHR mistrafficking into intracellular vesicles. Recent work in *Drosophila* revealed that Vps34 inactivation or pharmacological inhibition using the small molecule inhibitor, SAR405, caused alteration of cell polarity and disruption of epithelial architecture by relieving LKB1 inhibition and triggering JNK activation (14). A role of Vps34 in epithelial organization and polarity was also observed in 3D cultures of Caco-2 kidney cells (14). It would be interesting to analyze the activation states of LKB1 and JNK in Vps34^cKO^ thyroid, and, if modified, to cross Vps34^cKO^ with floxed LKB1 alleles.

The expression of the two SLC transporters, *Slc26a7* and *Slc26a4*, came out as a surprise in Vps34^cKO^. Decreased expression of Slc26a7 (4-fold) in Vps34^cKO^ may affect entry of iodine in the thyroid. However, it should be mentioned that this alternative basolateral transporter, SLC26A7, may only play an indirect role on iodine uptake (3). In addition, expression of the main transporter, *Nis*, is at least 16-fold more important than that of *Slc26a7*. This may explain why ^125^I uptake was not affected. Expression of apical *Slc26a4* (pendrin) in Vps34^cKO^ was even more dramatic with a 20-fold increase in expression. However, we do not think that this would have affected apical transport of iodine. Indeed, in control thyroids, expression level of *Slc26a4* were 60-fold less important than those of *anoctamin1* (6 Ct). This supports a more prominent role for *anoctamin1* in iodine apical transport, as demonstrated by Twyffels (36). These changes in expression of *Slc26a7* and *Slc26a4* could however impact on thyrocyte ion balance. Indeed, SLC26A7 and SLC26A4 transporters have opposite action on chloride ions at the basolateral and apical membranes, respectively. Decreased levels of SLC26A7 at the basolateral membrane may decrease the export of chloride out of the thyrocyte (3). On the contrary the increase of SLC26A4 at the apical membrane may increase the entry of chloride in the thyrocyte. Accumulation of chloride may in turn decrease intracellular pH and causes cellular stress in Vps34^cKO^.

The weak PAS staining in Vps34^cKO^ could be due to decreased exocytosis of Tg or increased Tg endocytosis. We indeed very often observed intracellular Tg in Vps34^cKO^. We suggest that the defective process is the endocytic route rather than exocytosis because we readily observed intracellular structures positive for iodinated Tg. This is rarely the case in control thyrocyte where I-Tg is rapidly proteolytically processed. In addition, our work also evidenced increased abundance and size of LAMP-1-positive late endosomes/lysosomes in Vps34^cKO^ thyrocytes, with accumulation of Tg and the autophagic marker, p62, indicative of a defective lysosomal trafficking and/or function. In addition to impaired Tg iodination, low T_4_ plasma levels thus also probably result from impaired endocytic transport of iodo-Tg to lysosomes and/or proteolytic excision of T_3_/T_4_ therein. Several studies have reported a role for Vps34 in endocytic trafficking to lysosomes (13, 15, 30, 37) and activation of lysosomal proteases (15). In our study on Vps34^cKO^ kidney PTCs, we also observed enlargement of lysosomes and their filling by undigested material, labelled for a variety of antigens (19). In Vps34^cKO^ thyrocytes, we also observed increased abundance and size of LAMP-1-positive compartments filled with Tg and I-Tg. Based on these and previous findings showing that Vps34 deletion leads to late endosome/lysosome enlargement and defective lysosomal function (37), we assume a defective proteolytic excision of T_3_/T_4_ from iodo-Tg, analogous to what we reported in cystinotic *Ctns^−/−^* mice (38). Cystinosis is a lysosomal storage disease due to deletion or inactivating mutations of the lysosomal cystine transporter, cystinosin (39). Defective cystinosin impacts on lysosomal proteolysis, presumably due to endolysosomal mislocalization, impaired pro-cathepsin maturation and altered luminal redox status; autophagic flux is also altered (40). Like cystinotic patients, *Ctns^−/−^* mice show compensated hypothyroidism with moderate increase of TSH, thyrocyte hyperplasia and proliferation, combined with progressive colloid depletion and evidence of increased endocytosis into colloid droplets. Iodo-thyroglobulin could be detected in *Ctns^−/−^* but not wild-type thyrocyte lysosomes, under identical labelling conditions, further indicating defective proteolysis.

It has recently become clear that Vps34 may act on lysosome positioning, which is crucial to autophagosome formation (41, 42). Similar to the kidney PTC defect, we observed an increased p62 signal in Vps34^cKO^ thyroids at P14. Accumulation of p62 and LC3b in Vps34^cKO^ indicates a block in the autophagy process. In addition, p62 accumulation also suggests activation of the antioxidant NRF2 pathway by competitive binding of p62 to KEAP1 (33). We indeed observed increased expression of two NRF2 target genes, namely the quinone reductase, *Nqo1*, and the thioredoxin reductase 1, *Txnrd1*. Considering the expected key role of Vps34 in macroautophagy, our data on mitophagy do not prove, but are compatible with the suggestion that this important homeostatic process is abrogated upon Vps34 deficiency.

A final observation deserving discussion is the occurrence of viable cells in the follicular lumens of Vps34^cKO^ thyroids. Our first hypothesis was that defective endocytic and autophagic routes to the lysosomes could impact thyrocyte homeostasis, thus causing cellular stress. Cellular stress, induced by defective autophagosome and lysosomal function, or by activation of stress kinase such as JNK, could explain shedding of thyrocytes cells into the colloidal space. However, luminal cells of Vps34^cKO^ thyroids were negative for E-cadherin and the thyrocyte-specific transcription factor, TTF-1. Furthermore, lineage tracing experiments demonstrated that these luminal cells had never expressed Pax8 and thus did not derive from thyrocytes. The second hypothesis was that the cellular stress and loss of tissue homeostasis would recruit macrophages. Indeed, we found that luminal cells were labelled for the conventional macrophage marker, F4/80, are proliferating and display high LAMP-1 signal. In addition, Tg and I-Tg colocalized with the LAMP-1 structures. This observation contributes to explain the colloid exhaustion observed in Vps34^cKO^ and could thus be responsible for the low levels of circulating T_4_. Although the signal(s) attracting macrophages to the colloid is(are) unknown, the loss of tight junction integrity might facilitate their invasion.

## Acknowledgements

This work was supported by grants from the Fonds pour la Recherche Scientifique (F.R.S-FNRS, # J.0076.18, Belgium), Université catholique de Louvain (Actions de Recherche concertées to CEP, ARC 15/20-065), and Fondation Roi Baudouin. SR and X-HL were supported by a grant DK15070 from the National Institutes of Health, USA. Work in the laboratory of BV was supported by grants from the MRC (G0700755) and BBSRC (BB/I007806/1; BB/M013278/1). The core facility for Imaging Cells and Tissues was also financed by National Lottery, Région bruxelloise, Région wallonne, Université catholique de Louvain and de Duve Institute. The authors thank Prof. C. Ris-Stalpers for iodo-thyroglobulin antibody, Abdelkadder El Kaddouri and Claude Massart for technical help. GG held a fellowship from the Fonds pour la formation à la Recherche dans l’Industrie et l’Agriculture (FRIA, Belgium), TW was postdoctoral researcher supported by UCLouvain, OD is supported by Télévie, VJ by the Cystinosis Research Foundation, CS is a UCLouvain teaching assistant, HGC was a postdoctoral researcher and CEP is a Senior Research Associate at F.R.S-FNRS.

## Author disclosures and contributions

BV is a consultant for Karus Therapeutics (Oxford, UK), iOnctura (Geneva, Switzerland) and Venthera (Palo Alto, US) and has received speaker fees from Gilead. The other authors have no competing financial interest. GG and TSW performed the initial morphological and molecular studies and analyzed the data. OD and CS performed all the other experiments and prepared the figures for the revision. VJ and AS participated in data collection and analysis. HGC was involved in supervision of GG. BB and BV provided the Vps34 mice, XHL and SR assayed T_4_ and TSH levels, and CM provided expertise with ^125^I experiments. PJC and CEP conceived, designed and supervised the project, and wrote the manuscript. All authors have approved the final version of the manuscript.

