## Supplemental material for "Vps34 PI 3-kinase controls thyroid hormone production by regulating thyroglobulin iodination, lysosomal proteolysis and tissue homeostasis"

GriecoSupplFig1

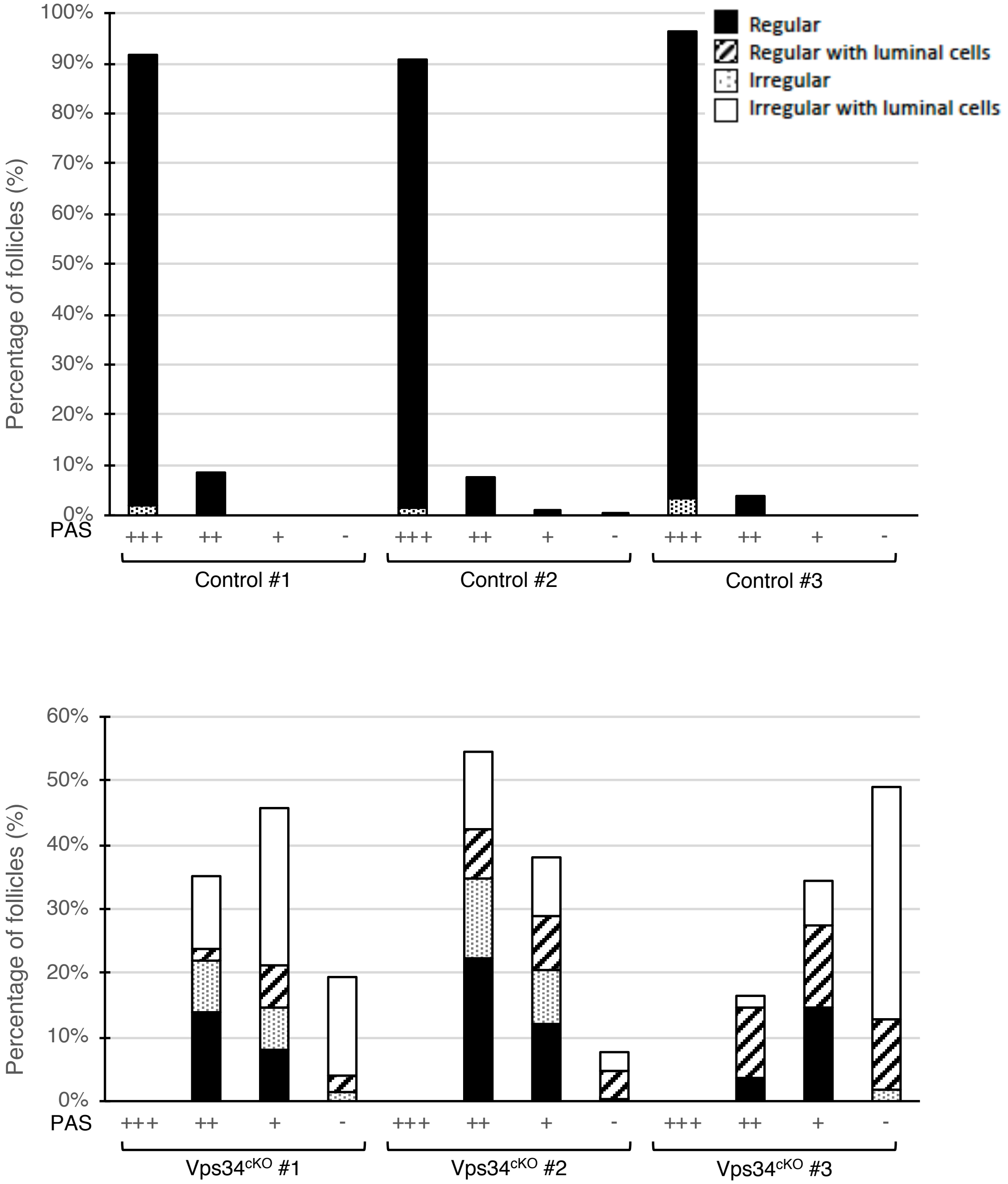


***Suppl Fig. 1. Histopathological evidence of colloid exhaustion*: quantification.** Follicles from three controls and three Vps34$^{cKO}$ sections stained with PAS were classified in four groups: regular follicle (black), regular follicle with luminal cells (dashed), irregular follicle (dotted) and irregular with luminal cells (white). In addition, each follicle group was assigned a PAS intensity: stronger than cartilage (PAS+++), similar to cartilage (PAS++), weaker than cartilage (PAS+) and no PAS staining (PAS-). In control, most follicles are regular and PAS++. In Vps34$^{cKO}$, the four type of follicles can be found and most follicles have low (PAS+) or no PAS staining.

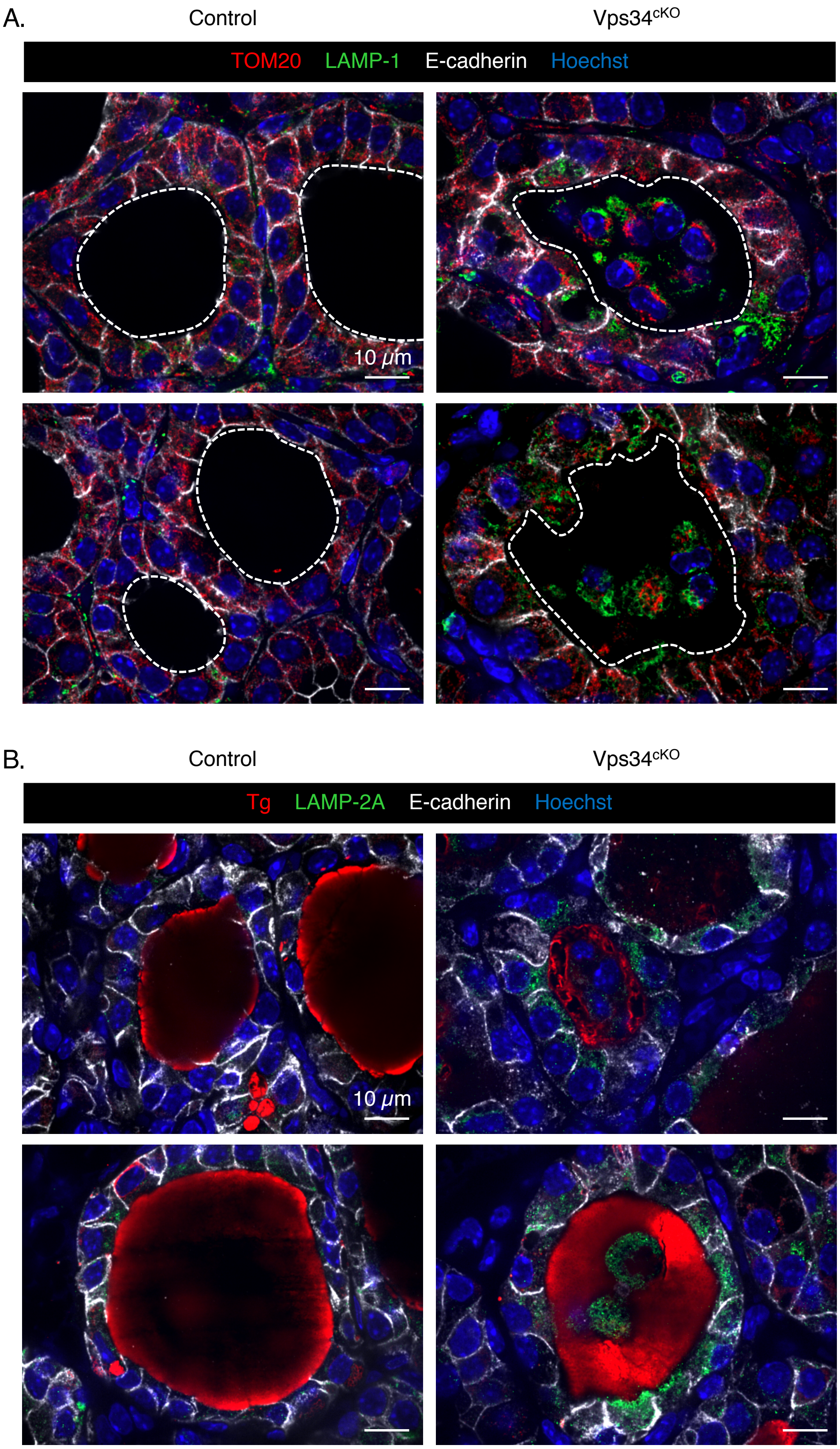


***Suppl Fig. 2. Macroautophagy and chaperone-mediated autophagy. A.*** Thyroid sections from control (left) and Vps34$^{cKO}$ (right) labeled for TOM20 (red), lysosomal LAMP-1 (green) and E-cadherin (white). Nuclei are labeled by Hoechst (presented in blue). In control and Vps34$^{cKO}$, TOM20 shows a perinuclear localization without co-localization with LAMP-1 (no yellow), even though LAMP-1 signal is more widespread. ***B.*** Thyroid sections from control (left) and Vps34$^{cKO}$ (right) labeled for thyroglobulin (red), LAMP-2A (green) and E-cadherin (white). Nuclei are labeled by Hoechst (shown in blue). In Vps34$^{cKO}$ sections, LAMP-2A signal is much stronger than in control thyroid.

*Supplementary Table I: Antibodies and use*

| Antibody | Supplier | Reference | Species | Dilution | Unmasking | Embedding |
|---|---|---|---|---|---|---|
| **Act. Casp. 3** | Cell signalling | 9661 | rabbit | 1/100 | + | Paraffin |
| **β-catenin** | BD Biosciences | 610154 | mouse IgG1 | 1/1000 | + | paraffin |
| **E-Cadherin** | BD Biosciences | 610182 | mouse IgG2a | 1/1000 | - or + | paraffin or gelatin |
| **Ezrin** | Thermo scientific | MS-661-P1 | mouse IgG1 | 1/400 | + | paraffin |
| **F4/80** | Cell signalling | 70076 | rabbit | 1/250 | + | paraffin |
| **I-Thyroglobulin** | Gift : Ris Stalpers | | mouse IgG1 | 1/100 | + | paraffin |
| **Ki67** | BD Biosciences | 556003 | mouse IgG1 | 1/250 | + | paraffin |
| **Laminin** | Sigma | L9393 | rabbit | 1/100 | + | paraffin |
| **LAMP-1** | Hybridoma Bank | 1D4B | rat (Mab) | 1/100 | - | paraffin or gelatin |
| **LAMP-2A** | Abcam | Ab18528 | rabbit | 1/200 | + | paraffin |
| **LC3B** | Cell signalling | 3868 | rabbit | 1/100 | + | paraffin |
| **$Na^+/K^+$-ATPase** | Hybridoma bank | α6F | mouse IgG2a | 1/400 | + | paraffin |
| **p62** | ARP | 03-GP62C | guinea pig | 1/400 | + | paraffin or gelatin |
| **TOM20** | Cell signalling | D8T4N | rabbit | 1/200 | + | paraffin |
| **Thyroglobulin** | DAKO | M0781 | mouse IgG1 | 1/500 | + | paraffin |
| **TTF1** | Agilent | M357501 | mouse IgG1 | 1/200 | + | paraffin |
| **YFP** | Abcam | Ab6673 | goat | 1/250 | + | Paraffin or gelatin |
| **ZO-1** | Invitrogen | 61-7300 | rabbit | 1/75 | + | paraffin |

*Supplementary Table II: Primers*

| gene | forward primer (5' → 3') | reverse primer (5' → 3') |
|---|---|---|
| *Ano1* | *GAGGCCAGTAGCCATCAGAG* | *CGTGAAGGAGATCACAAAGGC* |
| *β-actin* | TCCTGAGCGCAAGTACTCTGT | CTGATCCACATCTGCTGGAAG |
| *Duox* | TCCAGAAGGCGCTGAACAG | GCGACCAAAGTGGGTGATG |
| *Duoxa2* | CGTTAACATTACACTCCGAGGAACA | CAGAATGCCACCCACAGTGT |
| *Gpx2* | GCTTCCCTTGCAACCAGTTC | CTCCCCTTCTGGCCCTATGA |
| *Nis* | AGCAGGCTTAGCTGTATCCC | AGCCCCGTAGTAGAGATAGGAG |
| *Nqo1* | CGTCATTCTCTGGCCGATTCA | GGGGAAAAAGAAAGCTGCGT |
| *Nrf2* | GAATTCCTCCCAATTCAGCCG | GCTGCCTCCAGAGAGCTATT |
| *Rpl27* | GCCCTGGTGGCTGGAATTGACC | AAACTTGACCTTGGCCTCCCGC |
| *Slc26a4* | GCTCGCATTCGGGACTGTAA | CAGCAAACCTGCTTTGGCAT |
| *Slc26a7* | CTCAGTTCCCTGTCCAACGG | AGCGTGGAGACTCCTGTGTA |
| *Tg* | TGGGACGTGAAAGGGGAATGGTGC | GTGAGCTTTTGGAATGGCAGGCGA |
| *Tpo* | TGCCAACAGAAGCATGGGCAAC | GCACAAAGTTCCCATTGTCCAC |
| *Tshr* | CTGCGGGGCAAAGAGTGTGC | AGGGGAGCTCTGTCAAGGCA |
| *Txnrd1* | TACTGCATCAGCAGTGATGATC | CCATGTTCTTCCATGTGTTCAC |
| *Vps34 ex20-21* | CACACAGTATCCAGCACAGC | TGAGACTGGACCCTATGGCA |
| *Vps34 ex21-22* | TTGGAGTTGGAGACCGGCA | TCCTTGTTCAGCTTCATCGGAG |
| *Vps34 ex22-24* | GATGGTGGAAGGGATGGGTG | CGCTCTCGTCGATCAGACTC |
